# Pixel Quantum Efficiency Differences and Variance Stabilization for sCMOS Single Molecule Localization Microscopy Data Analysis

**DOI:** 10.1101/445452

**Authors:** Hazen P. Babcock, Fang Huang

## Abstract

Optimal analysis of single molecule localization microscopy (SMLM) data acquired with a CMOS camera requires compensation for single pixel differences in gain, offset and readout noise. For some CMOS cameras we found that it is also necessary to compensate for pixel differences in sensitivity or relative quantum efficiency (RQE). We present the modifications to the original sCMOS analysis algorithm necessary to correct for these RQE differences. We also discuss the use of the Anscombe transform (AT) for variance stabilization. Removing the variance dependence on the mean allows simpler least squares fitting approaches to achieve the Cramer-Rao bound on the mixed Poisson and Gaussian distributed data typically acquired with an sCMOS camera.

## Introduction

The high performance and relatively low cost of modern CMOS cameras makes their use attractive for SMLM. Recently they have started to displace EMCCD cameras in part because of their much greater bandwidth. A typical scientific CMOS camera can acquire 4M pixels at a frame rate of 100Hz, which is an order of magnitude faster than an EMCCD camera. This allows high throughput SMLM and also live cell SMLM with a large field of view^1–3^. This performance however comes with the downside of much greater pixel-to-pixel variability because unlike an EMCCD camera each pixel has it’s own read out amplifier. In order to get optimal results from the analysis of CMOS data it is necessary to compensate for these pixel-wise differences. The first demonstration of how to do this focused on handling differences in gain, offset and readout noise when fitting for the positions of the localizations^2^. Subsequent work included correction for RQE differences^4^.

As the largest pixel-to-pixel differences are found in the readout noise, offset and gain it is reasonable to focus on these properties. However, at least for some CMOS cameras it is also necessary to compensate for differences in the RQE of the pixels. In one camera that was tested in this work the standard deviation of the per pixel RQE was ∼5%. This difference is large enough to lead to systematic errors in localization fitting on the order of 1 nm, which could be a significant error source for some of the recent high precision SMLM based techniques^5,6^. Including the RQE differences in the fitting model removes this bias, restoring Cramer-Rao lower bound (CRLB) limited localization fitting performance.

Huang et al.^2^ demonstrated how to transform Gaussian distributed camera read noise to Poisson distributed noise. This transform let the authors achieve CRLB fitting performance using maximum-likelihood estimation (MLE) fitting of data with Poisson noise. However MLE fitting of data with Poisson noise, which is based on log-likelihood calculations, is generally considered to be slower and more complicated than least squares fitting^7,8^. A Poisson distribution can be transformed into a Gaussian distribution with high accuracy by several variance stabilization approaches. One popular and simple way to do this is to use the Anscombe transform^9,10^, which transforms a Poisson distribution into Gaussian distribution with a variance of 1.0. The transform is not perfect as the variance still depends on the mean for data with counts below approximately 4. Such low count values are however rarely encountered in SMLM images which often have an auto-fluorescence background on the order of 10 or more counts (in units of photo-electrons, e-). Once the data has been transformed in this fashion least squares fitting will achieve the CRLB, as least squares fitting is the maximum-likelihood estimator for data with Gaussian distributed noise.

## Results

Camera calibration data was acquired following the procedure in Huang et al.^2^. Movies that were 20k frames in length were acquired at 4 different illumination intensities and also in the dark. These movies were used to fit for the gain, offset, read noise and RQE of each pixel on the camera. First the offset and the read noise were determined by measuring the mean and the variance respectively of the dark movie. Then at each illumination intensity the mean and variance of the dark movie was subtracted from the movies measured mean and variance values. The gain was determined from the slope of best line to the corrected mean and variance values for each movie as well as a point at the origin, with equal weight given to every point. Finally the pixel values in the average image of calibration movie acquired at the highest intensity were converted to e- using the measured gain and offset. A smoothed version of the average image was created using the scipy.ndimage.uniform_filter() function with a size of 10 pixels^11^. The RQE for each pixel was calculated by dividing the averaged image by the smoothed image. The locally smoothed version of the average image was used instead of simply dividing by the mean value of the average image in the RQE measurement in order to reduce the effects of non-even uniform across the field of view.

An image of the camera gain for each pixel in a small region of the camera chip is shown in Fig. 1A. The standard deviation of the pixel gains is ∼10%. This is substantially larger than the error in the measurement, which is ∼1% (Supplementary Fig. S1). In Fig. 1B,C the gain and RQE of 10 different pixels from two different measurements are plotted. A visual comparison of these panels suggests that the gain and RQE are anti-correlated, pixels with a higher gain value have a lower RQE value. This anti-correlation is very clear in a 2D histogram of the gain versus RQE (Supplementary Fig. S2). Based on this data it appears that the camera manufacturer may have adjusted the pixel gains in order to compensate for quantum efficiency differences between the pixels. While this makes for a more visually pleasing picture it is problematic for quantitative analysis of the images. In particular many sCMOS localization fitting algorithms assume that every pixel has the same RQE. When this assumption is violated, especially in the non random fashion indicated by the vertical striping of the gain values here, the localization fits are systematically displaced^12^.

**Figure 1.**
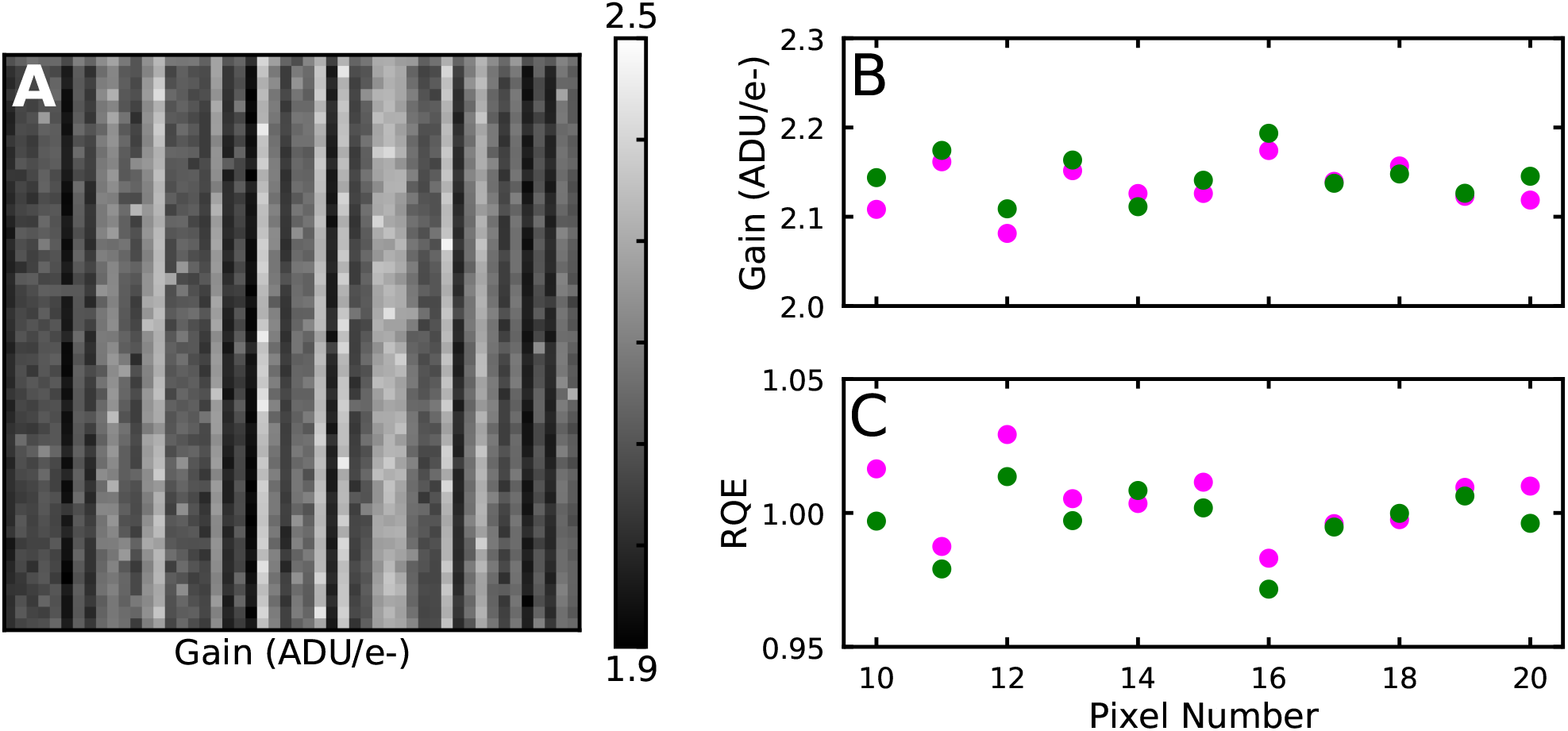
sCMOS camera calibration data from a ORCA-Flash4.0 (Hamamatsu photonics) camera. (a) The measured gain for each pixel in a small region of the sCMOS camera. (b) A graph of the measured gain at 10 different pixels. (c) A graph of the measured relative QE for the same 10 pixels. Magenta and green symbols are the values from two independent measurements.

Localization fitting was done using the Poisson MLE approach described in^13^. In this approach a variation of the Levenberg-Marquadt algorithm^14^ is used to minimize *χ*^2^ in the following equation:

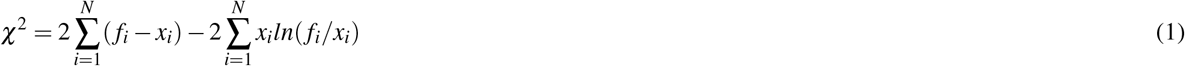

We tested several different options for the best fit image *f_i_* and the normalized camera image *x_i_*:

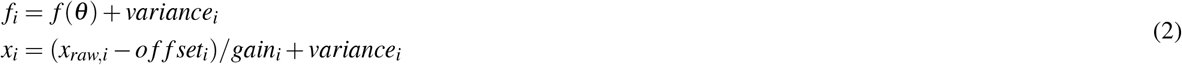

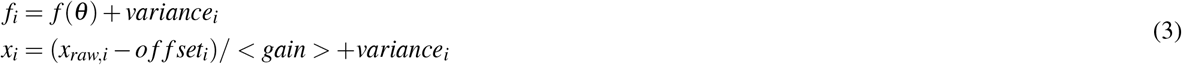

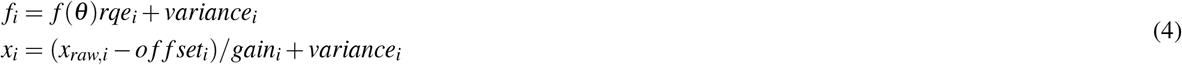

Where *f* (*θ*) is the fitting model and *θ* is the vector of fitting parameters.

To capture the magnitude of the systematic errors due to the RQE differences we used the measured calibration data to generated simulated movies of single emitters. An example simulated image is shown in Supplementary Fig. S3. When this data was fit with (2), which ignores RQE differences, the root mean square error (RMSE) between the measured localization position and the ground truth position is several nm greater than the CRLB (Fig. 2 “Uncorrected”). If the simulated data is instead fit with (3), where the gain value has been replaced with the average gain value, than the RMSE is substantially closer to the CRLB (Fig. 2 “Mean Gain Correction”). This is likely because this correction provides a gain value that is close to the actual gain value, but still preserves local smoothness in the normalized camera image *x_i_.* The “Mean Gain Correction” approach may be a good option unless very high fitting accuracy is required. In this case the use of (4) in the fitting model gives an RMSE that is very close the CRLB even at the highest emitter intensities tested (Fig. 2 “RQE Correction”).

**Figure 2.**
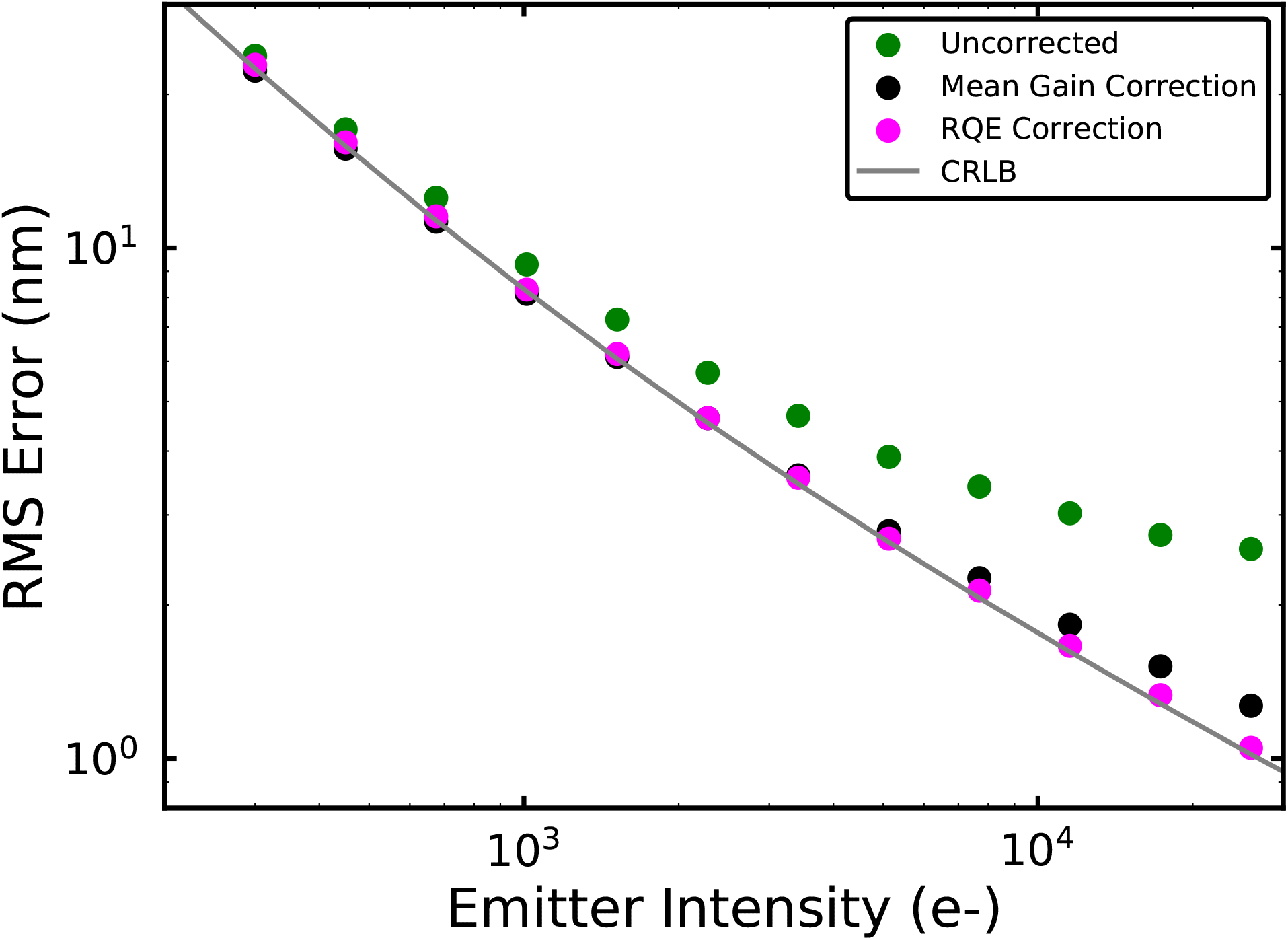
A graph of the effects of differences in relative QE on the accuracy of fitting. The measured sCMOS calibration data from a ORCA-Flash4.0 camera was used to simulate images of 12600 emitters for each e- value. The images included a constant background of 20e-. The images of the emitters were then fit using 3 different approaches. In the ‘Uncorrected’ approach the relative QE values were all set to 1.0 in the fitting. In the ‘Mean Gain Correction’ approach the relative QE values were set to 1.0 and all the pixels had the same gain value, the average of the pixel gain values. In the ‘RQE Correction’ approach the fit included the known relative QE and gain values for each pixel. CRLB is the Cramer-Rao lower bound calculated as described in Methods.

The above approaches differ from early work presented in Lin et al.^4^ which also includes a RQE correction. In Lin et al. the data is fit using a weighted least squares (WLS) approach, converted to the terminology used in this paper in equations 5 and 6. As the authors demonstrated, this approach achieves the CRLB for localizations with more than 300 photons, so from a practical perspective the differences between the two approaches is small. From a mathematical perspective however the important differences are that this work performs Poisson MLE fitting, and that this work includes the RQE term in the fit term *f_i_*. This is done in order to avoid distorting the Poisson distribution of *x_i_*, which is likely less of an issue for WLS fitting. If we had instead included the *rqe_i_* term in the equation for *x_i_* we would have ended up with something that is very close to the “Mean Gain Correction” approach (3). As discussed above, the *gain_i_* values for the camera used in this work have been adjusted such that 1 /(*rqe_i_*\**gain_i_*) is very close to 1/ < *gain* > for each pixel.

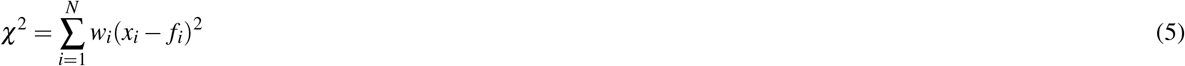

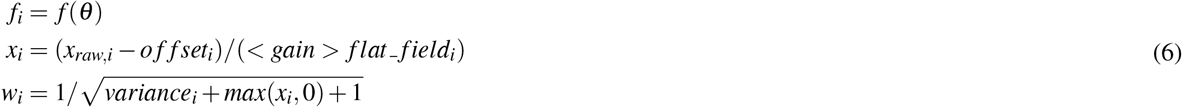

The Anscombe transform^9^ (7) is a simple way to transform Poisson distributed data into Gaussian distributed data. It is used in the SimpleSTORM algorithm^10^ for analysis of 2D SMLM data with a Gaussian PSF model. In this work we chose to apply the Anscombe transform to both the data and the fitting model. We opted for this approach because the Anscombe transform changes the shape of the PSF in a background dependent fashion. If the background is large relative to the signal, then the PSF after the Anscombe transform will be narrower than if the background is small relative to the signal. This is not such an issue with a Gaussian PSF model where it just introduces a background dependence to the *σ* of the Gaussian. It is much more of an issue for a cubic spline PSF model^15,16^, which would now need to include an extra dimension in the spline model to handle the PSF shape dependence on the background. An additional reason for our choice of this approach is that it required the least modification to other parts of our existing analysis pipeline.

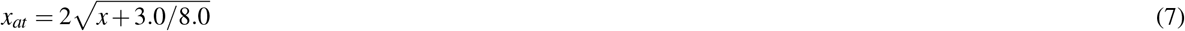

Poisson MLE fitting (PMLE) using equations 1, 4 versus Anscombe least squares (ALS) fitting using equations 8, 9 is compared in Fig. 3. The comparison was done on simulated sCMOS data with a fixed width 2D Gaussian fitting model 10. As shown in Fig. 3A the performance of these two approaches is essentially identical, and both achieve the CRLB. The ALS approach did not converge in significantly fewer iterations as might be expected for least squares fitting using the Levenberg-Marquadt algorithm. In order to make this comparison as fair as possible the convergence metric that we used was based on the delta values of a localizations best fit parameters. We considered the fit to a localization to have converged if the difference between the new and the old parameters after a single iteration of fitting parameters update was less than a fixed small value, for example 1.0e-3 pixels for the X and Y position. We also found that the convergence of both approaches was essentially identical when we used a variable width 2D Gaussian model (11) ((Supplementary Fig. S4).

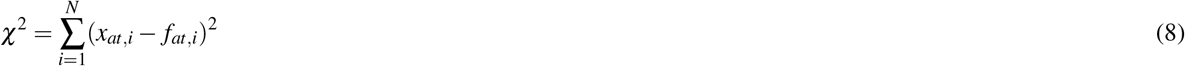

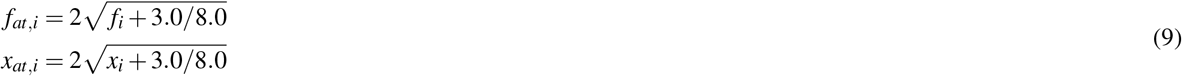

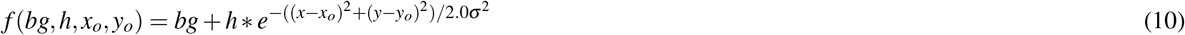

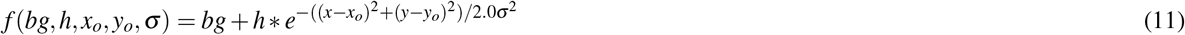

**Figure 3.**
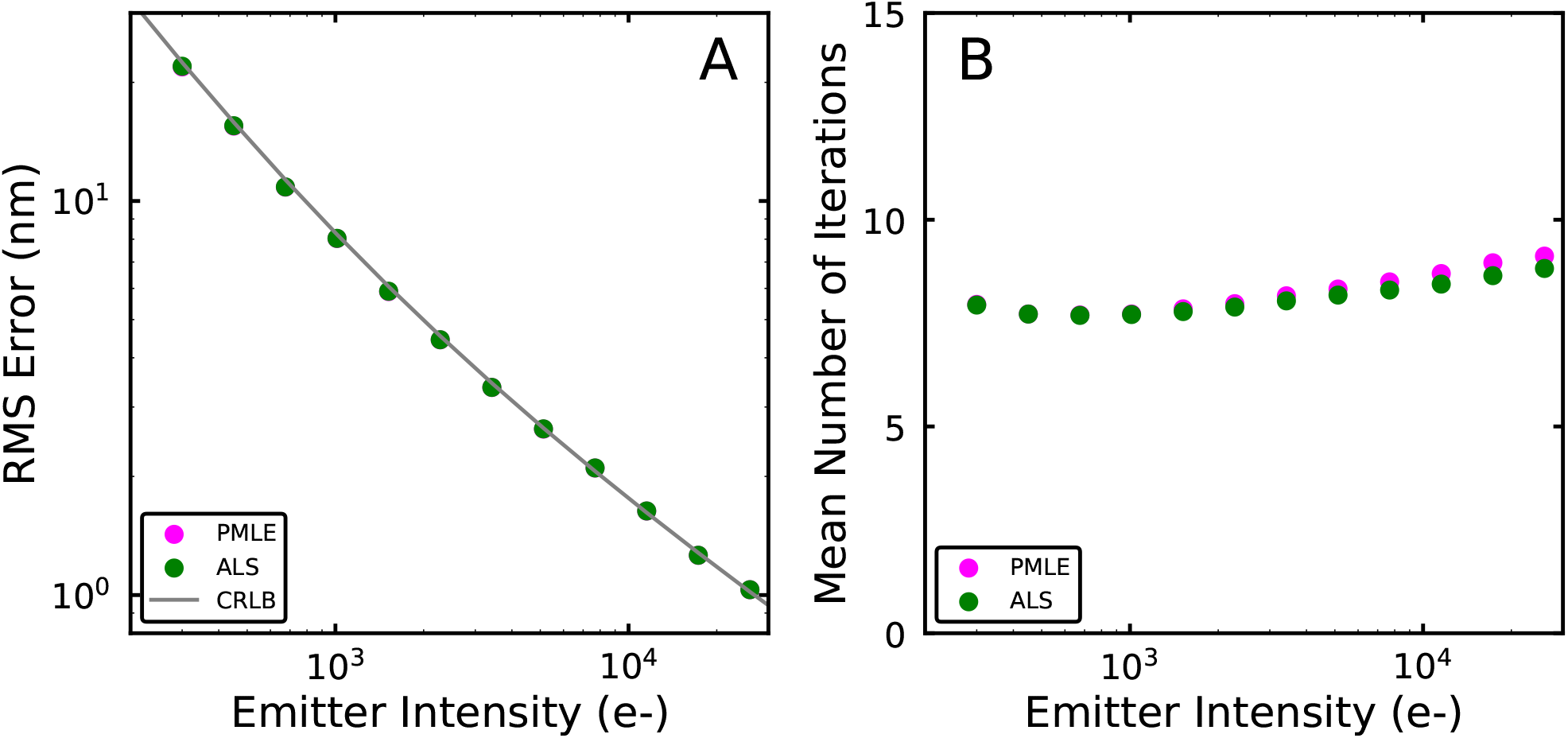
Graphs comparing the performance of Poisson MLE fitting (PMLE) versus least squares fitting of data whose variance has been stabilized using the Anscombe transform (ALS). The measured sCMOS calibration data from a ORCA-Flash4.0 was used to simulate images of 2520 emitters for each e- value. The images included a constant background of 20e-. The same images were then fit with fixed width Gaussians using the PMLE and the ALS fitting approaches. (A) A graph of the fitting accuracy of the two approaches. The CRLB line is the Cramer-Rao lower bound calculated as described in Methods. The performance of the two approaches is essentially identical so in most cases the PMLE and ALS points are indistinguishable. (B) The mean number of iterations per localization that each fitting approach took to converge.

In Fig. 3 the initial fit parameters were the ground truth values used in the simulations. It is possible that there was little difference in the convergence of the ALS versus the PMLE fitting approaches because the initial parameters were so close to the optimum. To test this as well as the robustness of the two approaches we explored the effect of displacing the starting x value from the ground truth value as shown in Fig. 4. Again there was little difference between the two approaches. Fig. 4A shows that the convergence of the two approaches is equally robust against poor x starting values. Fig. 4B shows that it takes an equal number of iterations for the fits that converged to converge regardless of the fitting approach. This was also unexpected as it is generally believed that least squares fitting is more robust than Poisson MLE fitting.

**Figure 4.**
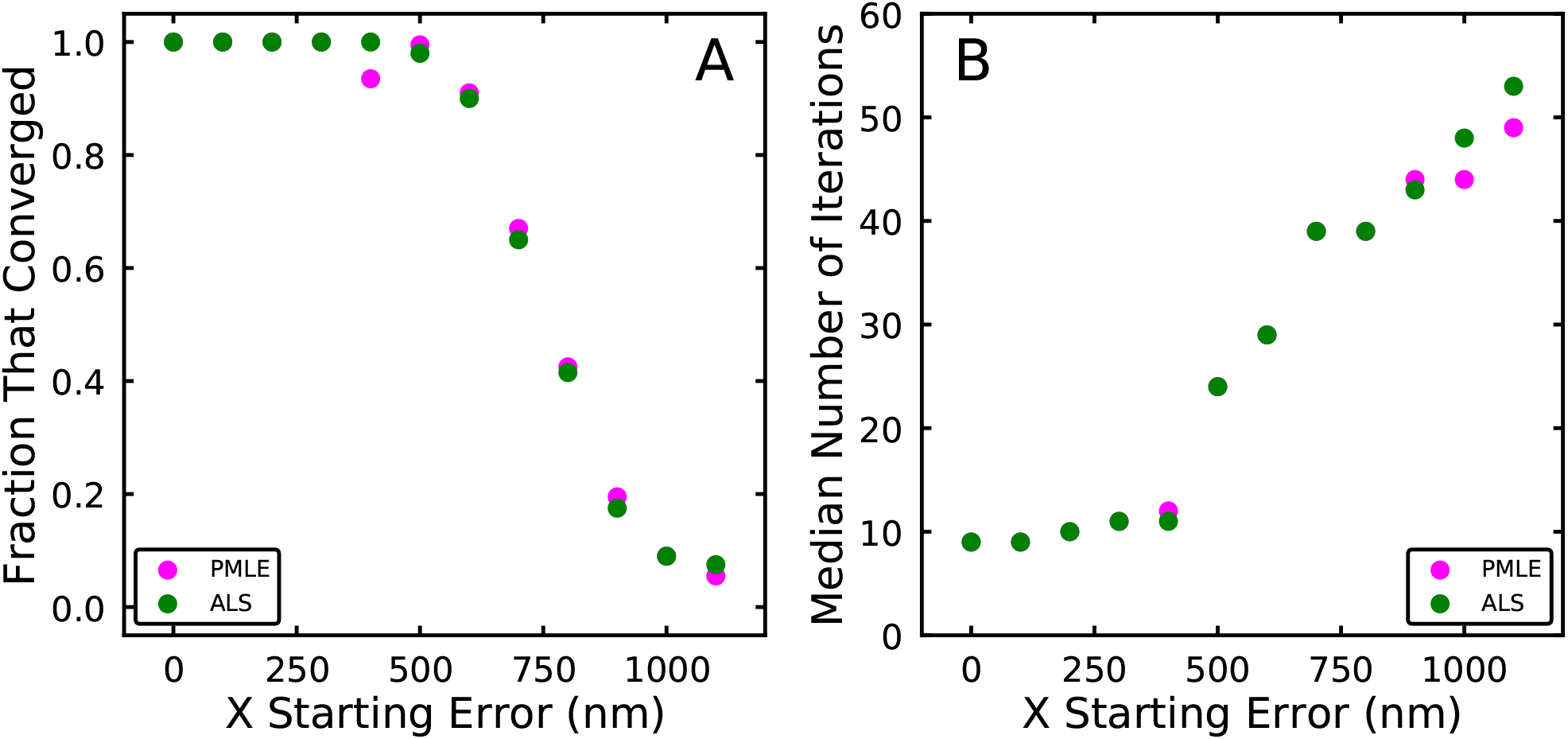
Graphs comparing the robustness of the Poisson MLE fitting (PMLE) and the least squares fitting of data whose variance has been stabilized using the Anscombe transform (ALS). The measured sCMOS calibration data from a ORCA-Flash4.0 was used to simulate images of 200 emitters for each x displacement value. The images included a constant background of 20e-. The same images were then fit with fixed width Gaussians using the PMLE and the ALS fitting approaches. (A) A graph of the fraction of fits that converged for each fitting approach as a function of the amount of error in the x position starting value. (B) The median number of iterations per localization that each fitting approach took to converge.

The most significant advantage of ALS that we found is that it is ∼80% faster than the PMLE fitting approach. In tests on a laptop with a i7-4510U CPU (Intel) ALS was able to perform 257k fitting iterations a second versus 143k fitting iterations a second for PMLE. Additional profiling indicates that this is primarily due to the fact that this CPU can perform the C language sqrt() function ∼10x faster than the log() function. The additional fitting speed however only results in a modest improvement of ∼5% in the overall speed of the analysis pipeline. This is due to the overhead of other parts of the pipeline such as image segmentation.

To verify that the ALS fitting approach also worked well on experimental data we used it to re-analyze an Alexa-647 labeled microtubule dataset acquired previously^17^. As shown in Fig. 5 the results of the ALS fitting approach are visually identical to those of PMLE fitting approach. Both approaches also identified almost exactly the same number of localizations, 8.32M for PMLE versus 8.30M for ALS.

**Figure 5.**
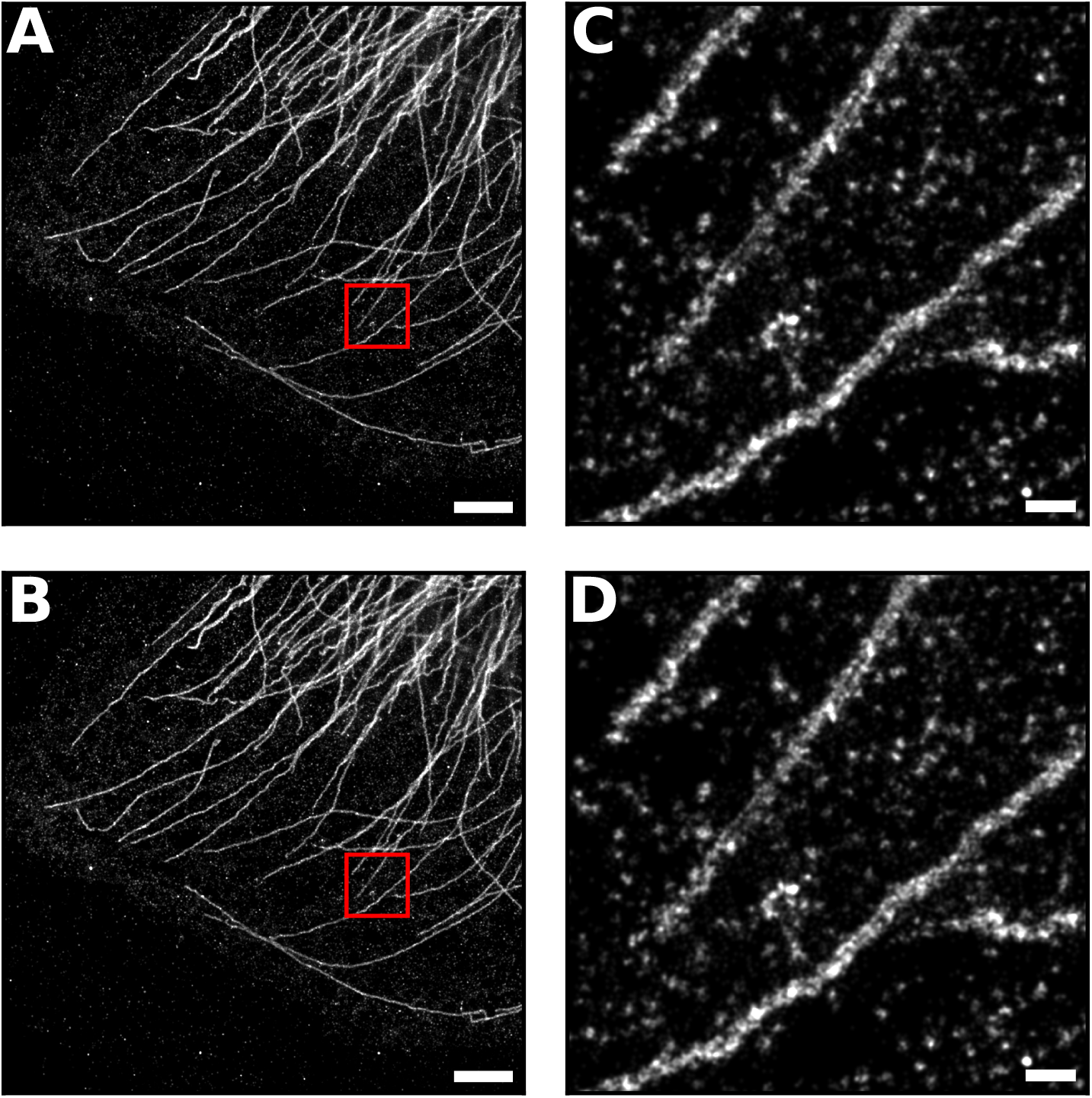
Super-resolution images of Alexa-647 labeled microtubules in U2OS cells analyzed with Poisson MLE fitting (PMLE) and least squares fitting with variance stabilization using the Anscombe transform (ALS). (A) A super-resolution image created from a STORM movie that was analyzed with PMLE. (B) A super-resolution inage creating by analyzing the same movie with the ALS. (C) Zoomed image of the red box in (A). (D) Zoomed in image of the red box in (B). Scale bars are 2um in A, B and 200nm in C, D.

## Discussion

In this work we presented an approach to correct for the significant pixel to pixel RQE differences encountered in some sCMOS cameras. This approach removes the bias in localization position caused by the RQE differences and restores CRLB fitting performance. We also discuss the use of the Anscombe transform for analyzing SMLM movies. With this transform CRLB fitting performance can be achieved using least-squares fitting. This will hopefully prove to be useful because least-squares fitting is simpler to implement. In addition there are a large number of off the shelf software packages for least squares fitting. It could also be useful when applying compressed sensing approaches to SMLM analysis as the most popular compressed sensing algorithms such as FISTA^18^ expect data with Gaussian distributed noise.

## Methods

### Microscope setup

The setup is based on an inverted microscope (TiU, Nikon) mounted on an optical table (RS2000, Newport). This microscope has a brightfield lamp and condenser (Ti-C-LWD 0.52, Nikon) which was used to provide spatially uniform and approximately constant illumination for camera calibration. The sCMOS camera (0RCA-Flash4.0 v2, Hamamatsu Photonics) was mounted directly onto the left port of the microscope with no additional magnification.

### Camera calibration

A 10x 0.3NA air immersion objective (CFI Plan Apo Lambda 10x 0.45NA, Nikon) was used without any sample mounted on the microscope to provide spatially uniform illumination of the camera chip. The stability of the intensity of the brightfield lamp was measured by taking the average intensity of each frame of the calibration movie. Contributions to the pixel variance due to fluctuations in the intensity of the brightfield lamp were ∼5% at the highest intensity used in calibration. Though small, these fluctuations were corrected for by subtracting the variance of the average intensity from the variance for each pixel.

### Cramer-Rao lower bound calculation

The Fisher information matrix is given by 12 where *θ* are the fitting parameters.

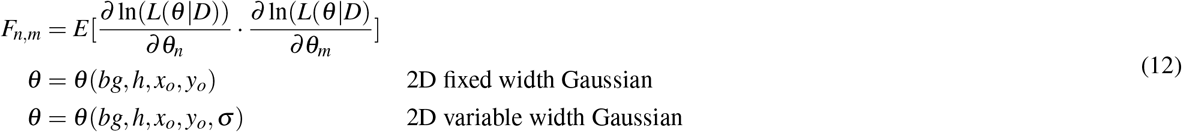

For a Poisson process with independent pixels (*i*) we can use the Stirling approximation to simplify 12 to 13. In this equation *μ_i_*(*θ*) is the fitting model.

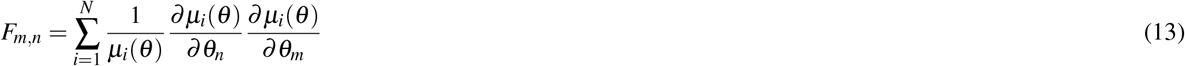

With the relative QE and sCMOS noise modifications *μ_i_*(*θ*) is given by 14

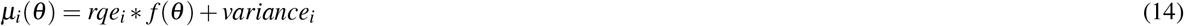

Finally substituting this into 13 gives 15.

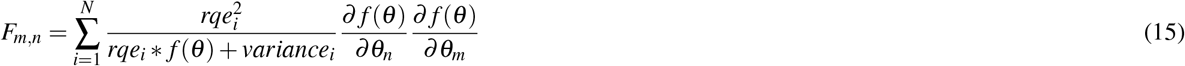

The Cramer-Rao bounds are then calculated from the inverted *F_m,n_* matrix.

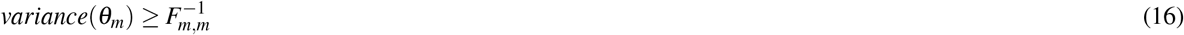

This calculation is slightly more complicated than the CRLB calculation in^7,19^ because the bounds now depend on the localizations position in the image. The CRLB values used in the figures are the average CRLB of all the simulated localization positions.

### Simulations and Analysis

All simulations and analysis were done using the open source storm-analysis project^20^. This project provides a mixture of Python and C language code that implements many of the common tasks in SMLM movie analysis. Jupyter^21^ notebooks that perform the simulations and create the figures are available in the supplementary material.

## Data Availability

Data in this paper is available by request.

## Acknowledgements

H.B. was supported by the Center for Advanced Imaging at Harvard University. F.H. is supported by grants from the NIH (R35 GM119785) andDARPA (D16AP00093).

## Author contributions statement

H.B. performed the simulations and experiments. H.B. and F.H. supervised the project. All authors contributed to writing the paper.

## Additional information

### Supplementary information

accompanies this paper at..

## Supplementary Material

**Supplementary Figure S1:**
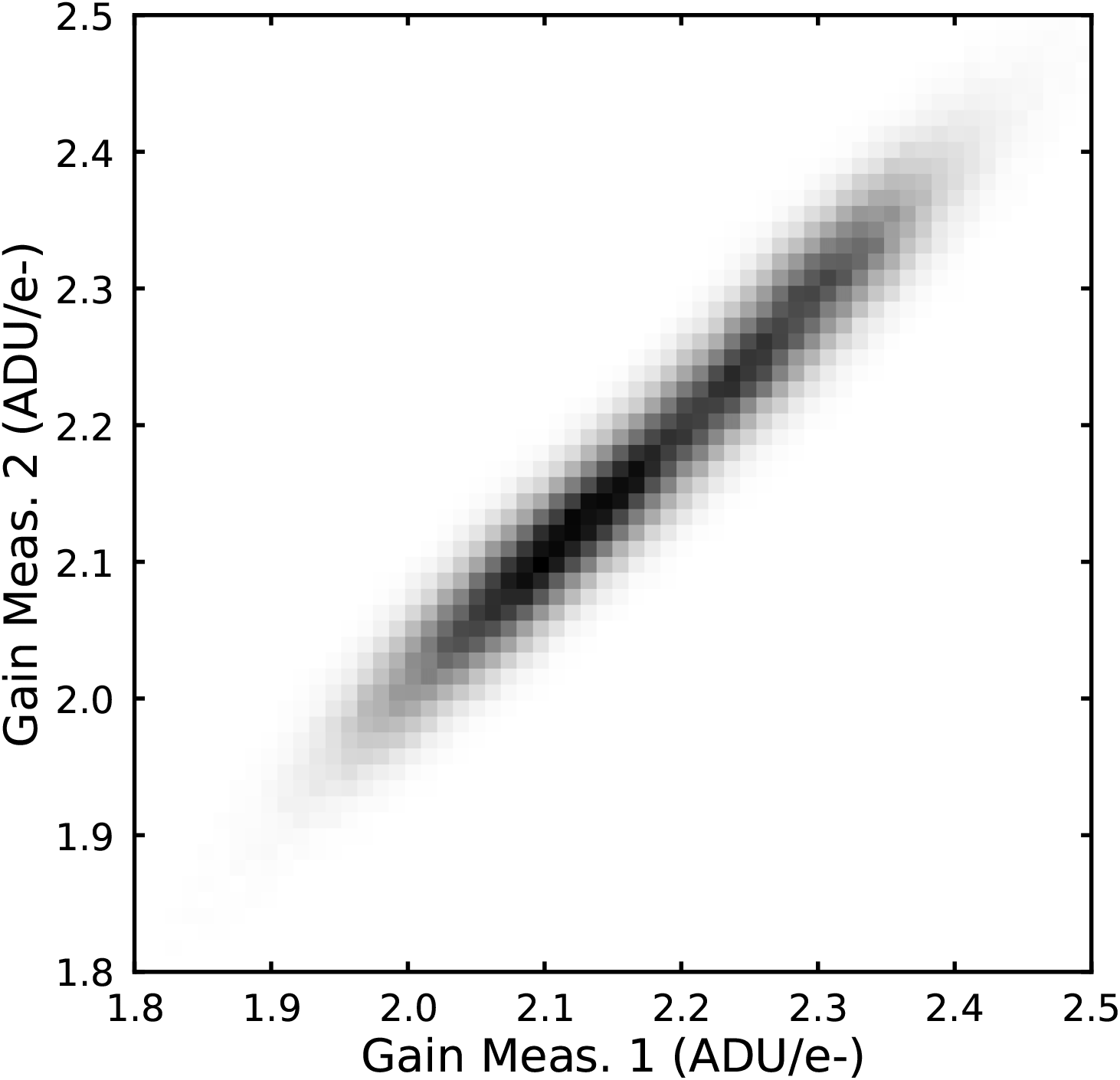
sCMOS Calibration Repeatability. A 2D histogram of the gain values measured in two different experiments to verify the repeatability of the camera calibration measurements.

**Supplementary Figure S2:**
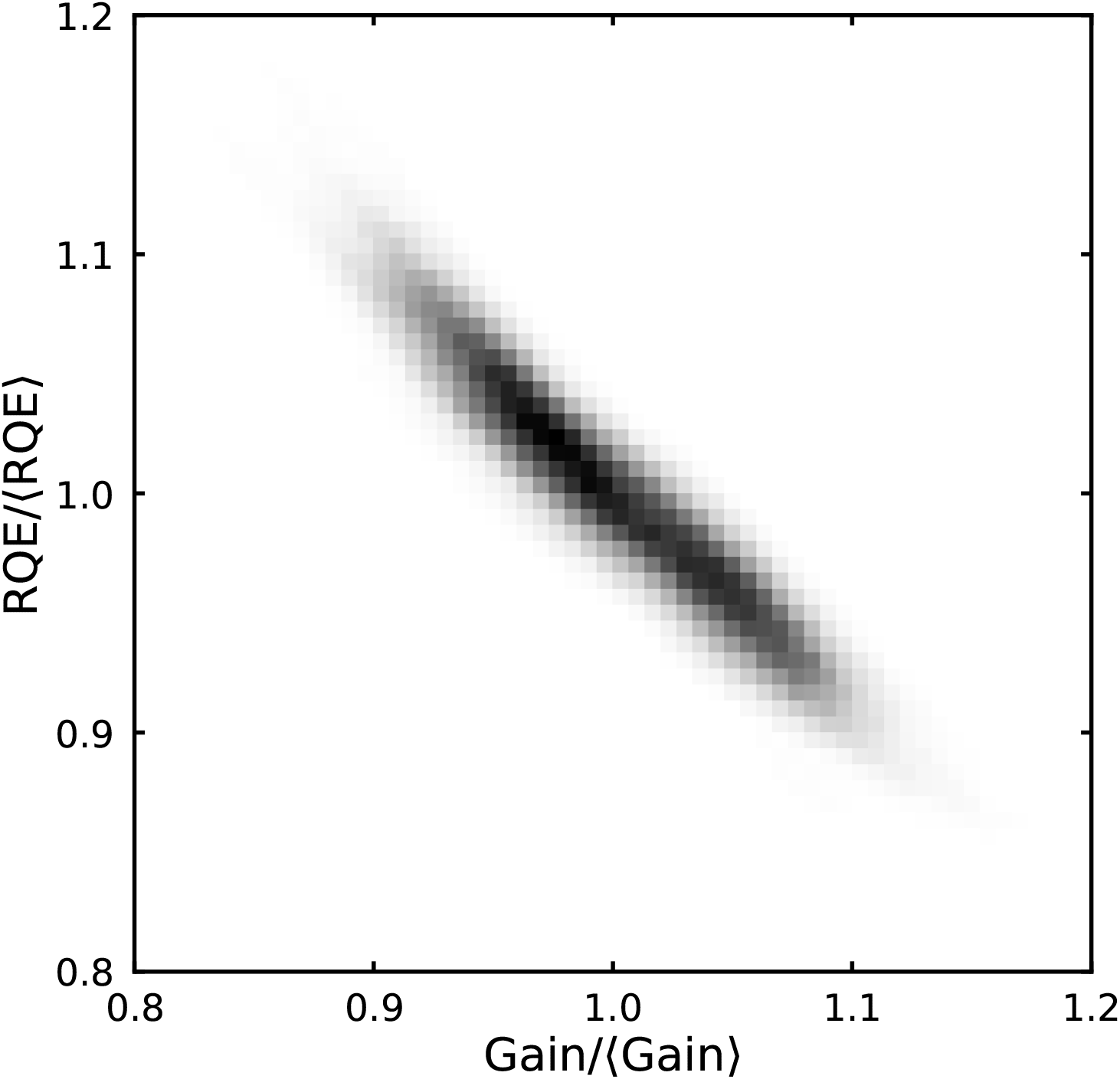
The Correlation between Gain and Relative QE. A 2D histogram of the normalized gain versus the normalized relative QE for each pixel. The strongly anticorrelated nature of these two values indicates that the gain is adjusted to compensate for differences in the relative QE of each pixel.

**Supplementary Figure S3:**
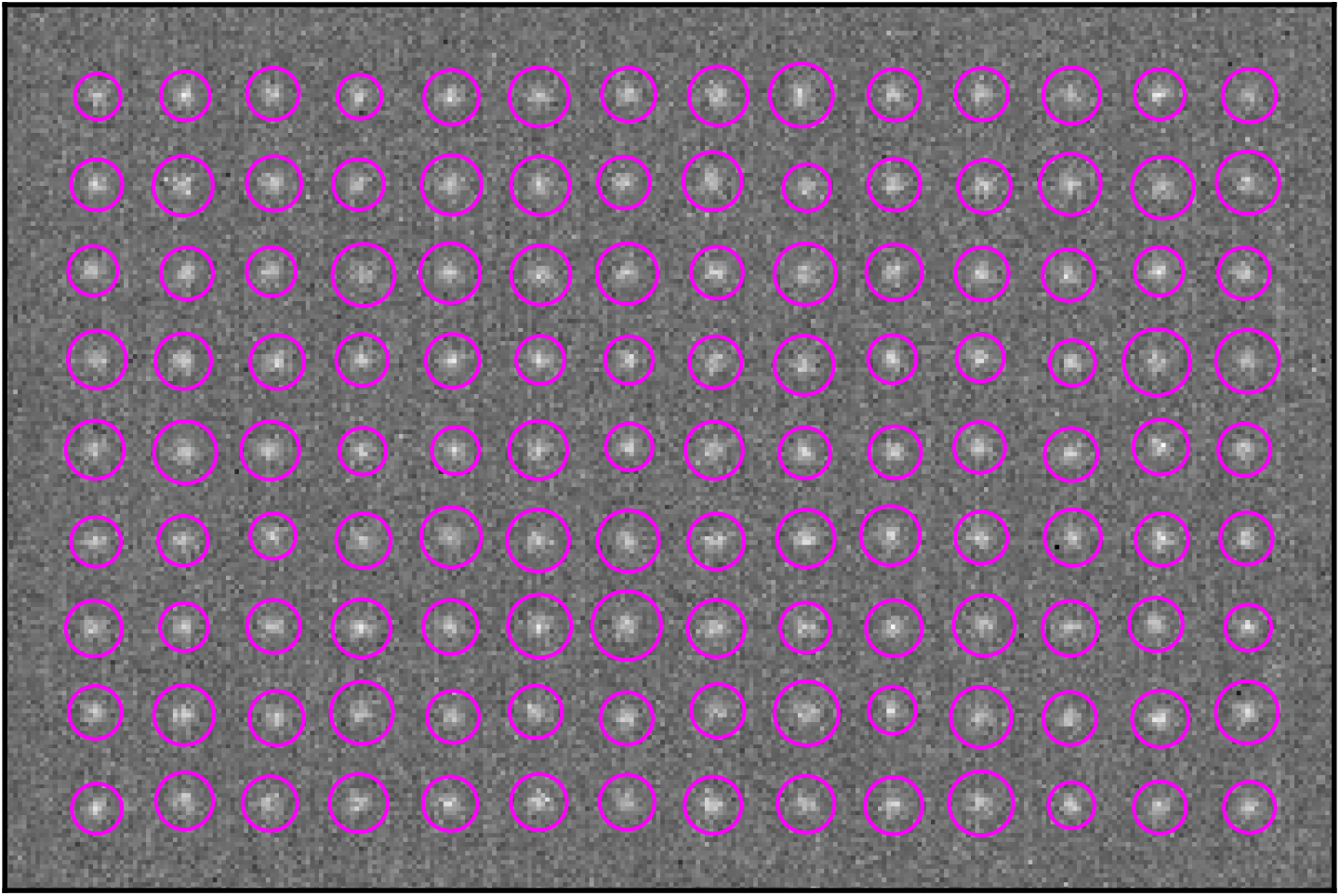
sCMOS Simulation Example. An example of the simulated images that were used to test the analysis approaches. The background in this image is 20e- and each localization has an integrated intensity of 450e-. The magenta circles are results from analysis of the image with the PMLE fitting approach.

**Supplementary Figure S4:**
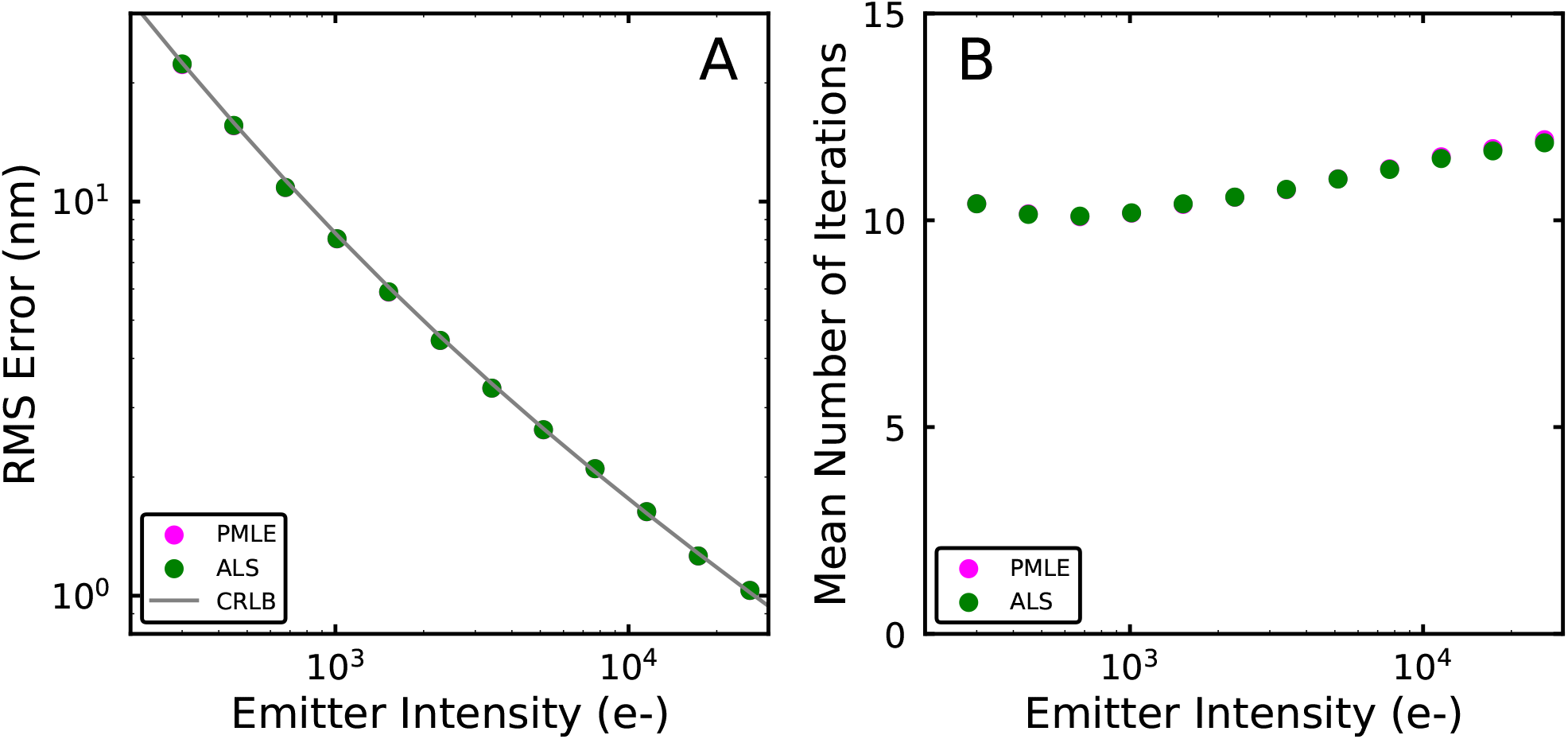
A Comparison of the Fitting Performance of PMLE versus ALS. Graphs comparing the performance of Poisson MLE fitting (PMLE) versus least squares fitting of data whose variance has been stabilized using the Anscombe transformation (ALS). The measured sCMOS calibration data from a ORCA-Flash4.0 was used to simulate images of 2520 emitters for each e- value. The images included a constant background of 20e-. The same images were then fit with variable width Gaussians using the PMLE and the ALS fitting approaches. (A) A graph of the fitting accuracy of the two approaches. The CRLB line is the Cramer-Rao lower bound calculated as described in the main text. The performance of the two approaches is essentially identical so in most cases the PMLE and ALS points are indistinguishable. (B) The mean number of iterations per localization that each fitting approach took to converge.

## References

1. Douglass, K. M., Sieben, C., Archetti, A., Lambert, A. & Manley, S. Super-resolution imaging of multiple cells by optimized flat-field epi-illumination. Nat. Photonics 10, 705–708 (2016).

2. Huang, F. et al. Video-rate nanoscopy enabled by sCMOS camera-specific single-molecule localization algorithms. Nat. Methods 10, 653–658 (2013).

3. Sigal, Y. M., Speer, C. M., Babcock, H. P. & Zhuang, X. Mapping synaptic input fields of neurons with super-resolution imaging. Cell 163, 493–505 (2015).

4. Lin, R., Clowsley, A. H., Jayasinghe, I. D., Baddeley, D. & Soeller, C. Algorithmic corrections for localization microscopy with scmos cameras - characterisation of a computationally efficient localization approach. Opt. Express 25, 11701–11716 (2017).

5. Balzarotti, F. et al. Nanometer resolution imaging and tracking of fluorescent molecules with minimal photon fluxes. Science 355, 606–612 (2017).

6. Jungmann, R. et al. Multiplexed 3D cellular super-resolution imaging with DNA-PAINT and exchange-PAINT. Nat. Methods 11, 313–318 (2014).

7. Mortensen, K. I., Churchman, L. S., Spudich, J. A. & Flyvbjerg, H. Optimized localization analysis for single-molecule tracking and super-resolution microscopy. Nat. Methods 7, 377–381 (2010).

8. Small, A. & Stahlheber, S. Fluorophore localization algorithms for super-resolution microscopy. Nat. Methods 11, 267–279 (2014).

9. Anscombe, F. J. The transformation of poisson, binomial and negative-binomial data. Biometrika 35, 246–254 (1948).

10. Köthe, U., Herrmannsdorfer, F., Kats, I. & Hamprecht, F. A. Simplestorm: a fast, self-calibrating reconstruction algorithm for localization microscopy. Histochem. Cell Biol. 141, 613–627 (2014).

11. Jones, E., Oliphant, T., Peterson, P. et al. SciPy: Open source scientific tools for Python (2001-). [Online; accessed 2018-07-26].

12. Pertsinidis, A., Zhang, Y. & Chu, S. Subnanometre single-molecule localization, registration and distance measurements. nature 466, 647–651 (2010).

13. Laurence, T. A. & Chromy, B. A. Efficient maximum likelihood estimator fitting of histograms. Nat. Methods 7, 338–339 (2010).

14. Marquardt, D. W. An algorithm for least-squares estimation of nonlinear parameters. J. Soc. for Ind. Appl. Math. 11, 431–441 (1963).

15. Babcock, H. P. & Zhuang, X. Analyzing single molecule localization microscopy data using cubic splines. Sci. Reports 7 (2017).

16. Li, Y. et al. Real-time 3d single-molecule localization using experimental point spread functions. Nat. Methods 15, 367–369 (2018).

17. Babcock, H. P. Multiplane and spectrally-resolved single molecule localization microscopy with industrial grade CMOS cameras. Sci. Reports 8 (2018).

18. Beck, A. & Teboulle, M. A fast iterative shrinkage-thresholding algorithm for linear inverse problems. SIAM J. on Imaging Sci. 2, 183–202 (2009).

19. Ober, R. J., Ram, S. & Ward, E. S. Localization accuracy in single-molecule microscopy. Biophys. J. 86, 1185–1200 (2004).

20. Storm-analysis, storm movie analysis code (2017). [Online; accessed 2018-07-26].

21. Project jupyter (2018). [Online; accessed 2018-07-26].

